# Acod1 Expression in Cancer Cells Promotes Immune Evasion through the Generation of Inhibitory Peptides

**DOI:** 10.1101/2023.09.14.557799

**Authors:** James H. Schofield, Joseph Longo, Ryan D. Sheldon, Emma Albano, Mark A. Hawk, Sean Murphy, Loan Duong, Sharif Rahmy, Xin Lu, Russell G. Jones, Zachary T. Schafer

## Abstract

Targeting PD-1 is an important component of many immune checkpoint blockade (ICB) therapeutic approaches. However, ICB is not an efficacious strategy in a variety of cancer types, in part due to immunosuppressive metabolites in the tumor microenvironment (TME). Here, we find that αPD-1-resistant cancer cells produce abundant itaconate (ITA) due to enhanced levels of aconitate decarboxylase (Acod1). Acod1 has an important role in the resistance to αPD-1, as decreasing Acod1 levels in αPD-1 resistant cancer cells can sensitize tumors to αPD-1 therapy. Mechanistically, cancer cells with high Acod1 inhibit the proliferation of naïve CD8^+^ T cells through the secretion of inhibitory factors. Surprisingly, inhibition of CD8^+^ T cell proliferation is not dependent on secretion of ITA, but is instead a consequence of the release of small inhibitory peptides. Our study suggests that strategies to counter the activity of Acod1 in cancer cells may sensitize tumors to ICB therapy.

## Introduction

The exploitation of the immune system to target cancer using the immune checkpoint blockade (ICB) strategy has revolutionized cancer treatment in recent years^1–3^. More specifically, antibodies targeting programmed cell death protein 1 (PD-1) can dampen the natural resistance of cytotoxic CD8^+^ T cells towards host cells and lead to increased recognition and destruction of cancerous cells^4^. Use of this strategy has shown strong efficacy in melanoma and non-small cell lung cancer patients^5–7^. Despite these exciting findings, many cancers are relatively insensitive (or entirely resistant) to αPD-1 (and other ICB) therapy^8, 9^. For instance, αPD-1 therapies in castration-resistant prostate cancer (CRPC) have largely failed to demonstrate durable clinical response^10,11^. As such, there remains a significant need to better understand the molecular mechanisms that underlie the sensitivity (or insensitivity) of cancer cells to ICB therapy.

The tumor microenvironment (TME) has emerged as a key determinant of sensitivity to αPD-1 ICB therapy^12^. In particular, extracellular metabolites present in the TME can often function to drive the immune response to a variety of malignancies^13–15^. For example, depletion of immunosuppressive metabolites within the TME reduces tumor size through enhancing the anti-tumor activity of cytotoxic CD8^+^ T cells^16, 17^. One metabolite that has recently garnered significant interest for its capacity to impact the immune response is itaconate (ITA). ITA is derived from the tricarboxylic acid (TCA) cycle through the enzymatic action of *cis*-aconitate decarboxylase 1 (Acod1, also known as Irg1) which catalyzes the decarboxylation of *cis*-aconitate to form ITA^18^. ITA has putative anti-inflammatory properties and has been revealed by multiple studies to have a significant role in regulating macrophage polarization^19–25^. Relatedly, *Acod1* expression appears to be upregulated under inflammatory stress in macrophages, monocytes, and dendritic cells^26, 27^. More recently, ITA has been discovered to have some capacity to regulate additional cell types, including neutrophils^28^, T cells^29, 30^, and epithelial cells^31^. Despite these important findings, the underlying mechanisms by which *Acod1* expression and ITA production alter behavior and function in other cell types remain inadequately characterized.

Here, we report that ICB-resistant cancer cells produce high amounts of ITA due to a significant elevation in Acod1 levels. This increase in Acod1 has a significant impact on ICB resistance as previously insensitive tumors, engineered to be deficient in Acod1, regain sensitivity to αPD-1 therapy. Mechanistically, cancer cells with high levels of Acod1 release factors into the extracellular space that block the activation and proliferation of CD8^+^ T cells. Surprisingly, the inhibition of CD8^+^ T cell proliferation is not due to ITA secretion by Acod1 high cancer cells. Instead, these cells produce small peptides that have the capacity to block CD8^+^ T cell proliferation. Taken together, our data suggest that strategies aimed at countering Acod1-mediated generation of small peptides in cancer cells may be efficacious in sensitizing tumors to ICB therapy.

## Results

### ITA is produced at high levels by ICB-resistant prostate cancer cells

In order to investigate links between changes in cancer cell metabolism and immune modulation, we utilized cell lines generated from a Probasin-driven *PB-Cre^+^ Smad4^L/L^/Pten^L/L^/Tp53^L/L^* mouse model of castration-resistant prostate cancer (CRPC)^32^. Tumors from these mice respond to αPD-1 immune checkpoint blockade (ICB) monotherapy and were used to derive a cell line for use in subsequent analyses (PPS-6239)^33^. To generate ICB-resistant variants of PPS-6239 cells, these cells were injected into syngeneic hosts, allowed to form tumors, treated with αPD-1, and were isolated from rare tumors that were not responsive to ICB monotherapy (Fig. S1). The resulting cells, termed PPS-PD1R, were resistant to αPD-1 monotherapy after injection into syngeneic hosts^33^. For the sake of simplicity, we will hereafter refer to the PPS-6239 cells as PRs (for **<u>pr</u>**ostate **s**ensitive) and the PPS-PD1R cells as PRp (for **<u>pr</u>**ostate **<u>P</u>**D1 resistant).

Having established these isogenic cell lines, we first sought to characterize metabolic changes associated with differing sensitivity to αPD-1 ICB therapy. We began this analysis by assessing mitochondrial metabolism in the PRs and PRp cell lines. Motivated by our recent findings demonstrating that detachment from extracellular matrix (ECM) can trigger mitophagy (the degradation of mitochondria in autophagosomes) in a fashion that causes cell death^34–36^, we assessed mitochondrial abundance in ECM-detached conditions. ICB-sensitive PRs cells indeed display lower mitochondrial mass during ECM-detachment as evidenced by the reduction in the MitoTracker signal (Fig. 1a-c) and diminished levels of mitochondrial proteins (Fig. S2). In contrast, ICB-insensitive PRp cells maintain mitochondrial mass in ECM-detached conditions (Fig. 1a-c, Fig. S2), suggesting that they are better able to successfully adapt to ECM-detachment in a fashion that promotes tumor progression and/or metastasis^37, 38^.

**Figure 1:**
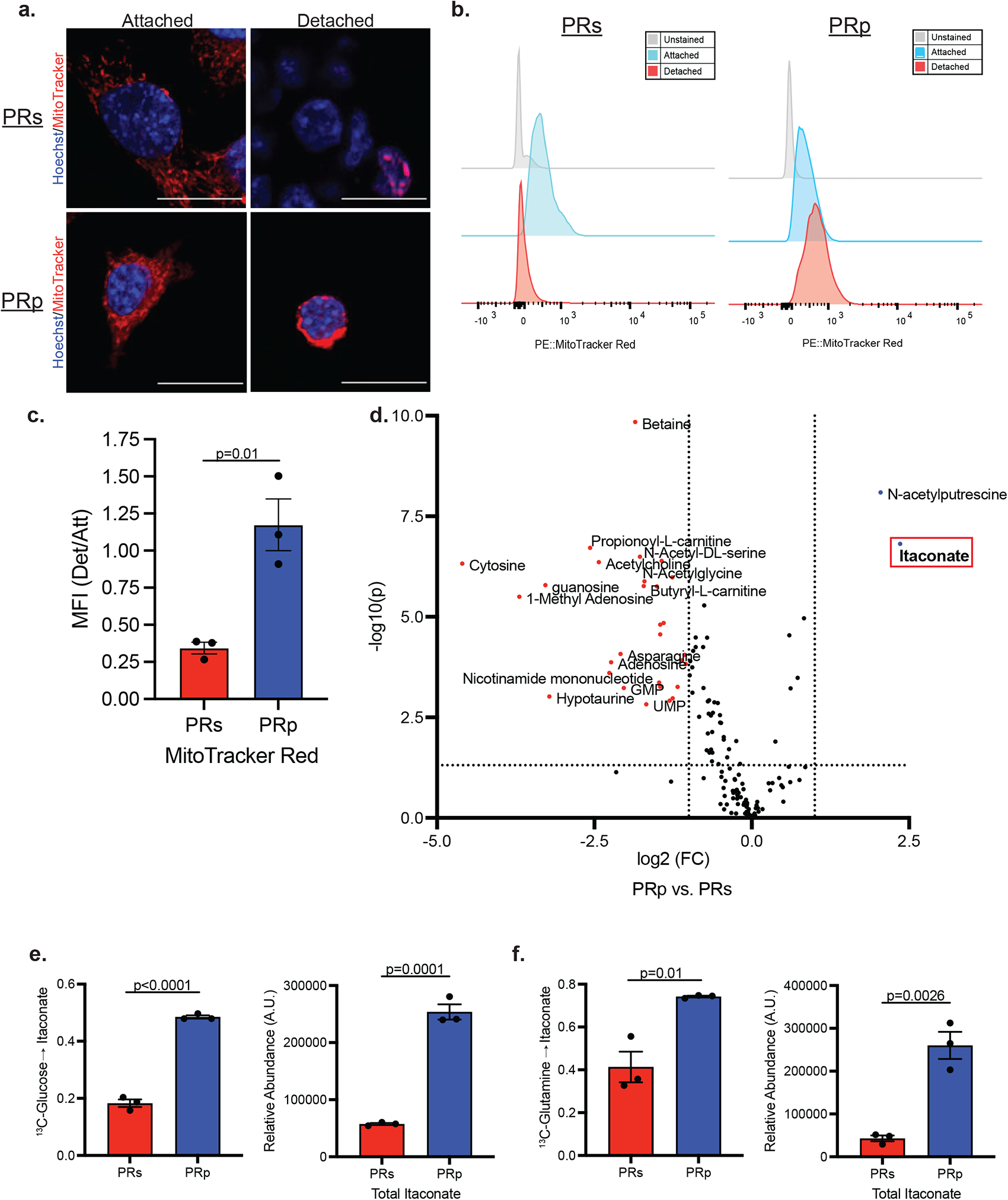
Itaconate is produced at high levels in ICB-resistant prostate cancer cells. PRs or PRp cells were grown in the indicated conditions for 24 h. **a.** Representative confocal images of cells stained with MitoTracker Red (red) and Hoechst 33342 (blue). Scale bars, 20 μm. **b, c.** Flow cytometry of cells stained with MitoTracker Red. Representative histograms (**b**) and mean fluorescence intensity (MFI) expressed relative to the corresponding attached (Att) culture condition (**c**). *n* = 3 independent biological samples. **d.** Volcano plot for intracellular metabolites in PRs and PRp cells after 24 h of detachment (Det) *P* value <0.05 and fold-change (FC) > 1.2 used for cutoffs. **e.** Fractional enrichment of labeled ITA (left) and total pool (right) from [U-^13^C]glucose tracing. **f**. Fractional enrichment of labeled ITA (left) and total pool (right) from [U-^13^C]glutamine tracing. Labeling done for 4 h in detached cells. Graphs represent data collected from a minimum of 3 biological replicates. *P* values are calculated by two-tailed Student’s *t* test. Data are mean ± SEM.

Given these results, we reasoned that maintenance of mitochondrial abundance during ECM-detachment may cause ICB-insensitive cells to utilize distinct metabolic pathways. To better understand metabolic differences between the ICB-insensitive and-sensitive cells, we conducted intracellular metabolomic profiling of PRs and PRp cells cultured in ECM-detached conditions. Surprisingly, itaconate (ITA) was the most significantly elevated metabolite in the PRp cells compared to the PRs cells (Fig. 1d). Given the abundance of ITA produced in PRp cells, we next sought to determine if the elevated levels of ITA was synthesized by flux originating from glucose or glutamine, two carbon sources widely utilized by cancer cells^39^. Indeed, labeling cells with [U-^13^C]glucose (Fig. 1e) or [U-^13^C]glutamine (Fig. 1f) revealed that PRp cells direct carbon flux from both nutrient sources into elevated production of ITA at a level that is significantly higher than PRs cells.

### Acod1 levels are elevated in ICB-resistant cells

As mentioned previously, the majority of studies on ITA have been conducted on cells of the monocyte-macrophage lineage, and we were therefore surprised by its detection in prostate cancer cells^40–42^. We next sought to investigate if the changes in ITA in PRp cells are associated with changes in Acod1, the enzyme responsible for ITA production (see diagram in Fig. 2a)^18^. Indeed, we found that Acod1 protein levels are elevated in the PRp cells when compared to PRs cells and that ECM-detachment promotes an increase in Acod1 levels in both cell lines (Fig. 2b). Furthermore, expression of *Acod1* transcript mirrored the protein data as mRNA levels are higher in PRp cells compared to PRs cells and similarly stimulated by ECM-detachment (Fig. 2c). These findings are not limited to the PRs and PRp cell lines, as we found that PC3 cells, a well-established human prostate cancer cell line, have discernable levels of ACOD1 protein that is increased as a consequence of ECM-detachment (Fig. S3a). We also expanded our analysis of ACOD1 protein levels into other cancer cell lines of diverse origin, and we found robust levels of ACOD1 protein in the GCT (derived from a fibrous histiocytoma) and MDA-MB-436 (derived from a pleural effusion in a triple negative breast cancer patient) cell lines (Fig. S3b).

**Figure 2:**
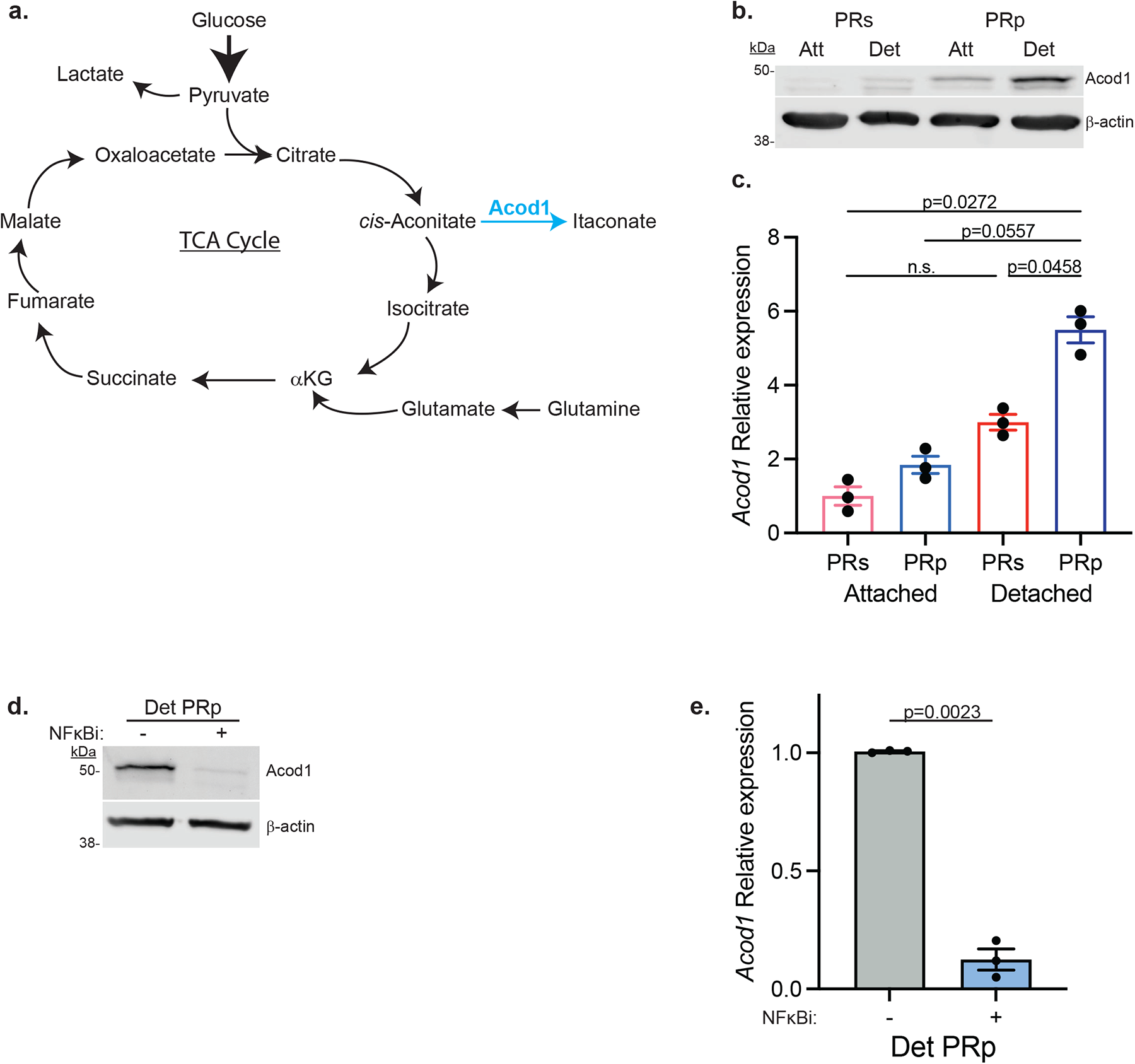
ACOD1 levels are elevated in ICB-resistant cells. **a**. Schematic of ITA generation by *cis*-aconitate decarboxylase (Acod1). **b**,**c**. PRs or PRp cells were grown in the indicated conditions for 24 h. **b,** Lysates were collected and immunoblotted as noted. **c**, Gene expression of *Acod1* by quantitative real-time PCR calculated as fold-change relative to Att PRs. **d**,**e**. PRp cells were grown in Det for 9h in the presence of NFκB inhibitor BAY1170-82 (2.5 μM). **d**, Lysates were collected and immunoblotted as noted. **e**, Gene expression of *Acod1* by quantitative real-time PCR. Graphs represent data collected from a minimum of 3 biological replicates and all western blotting experiments were independently repeated a minimum of three times with similar results. *P* values are calculated by one-way ANOVA followed by Tukey’s test in (**c**); two-tailed Student’s *t* test in (**e**). Data are mean ± SEM.

We next examined how Acod1 levels are regulated in prostate cancer cells. Previous studies have determined that diminished integrin function in epithelial cells during separation from the ECM activates the NFκB signaling pathway^34, 43^. Relatedly, *Acod1* expression has previously been found to be regulated by NFκB signaling^44^. Thus, we assessed the contribution of NFκB signaling to the increase in Acod1 levels observed in PRp cells. Indeed, treatment of detached PRp cells with BAY1170-82, an NFκB inhibitor, caused a significant downregulation of both Acod1 protein (Fig. 2d) and *Acod1* mRNA expression (Fig. 2e). We therefore conclude that NFκB signaling contributes to the ECM-detachment-induced Acod1 increase in prostate cancer cells.

### Acod1 dictates sensitivity to αPD-1 immune checkpoint blockade *in vivo*

Given the elevation in Acod1 levels in the PRp cells, we were interested in whether Acod1 contributes to the insensitivity of PRp tumors to αPD-1 monotherapy (see schematic in Figure 3a). As such, we engineered PRp cells to be deficient in Acod1 (Figure 3b) and then inoculated control (PRp Scr) or Acod1 knockout (PRp Acod1 KO) cells subcutaneously into male C57BL/6 mice. After 11 days of tumor growth, we subjected these mice to αPD-1 monotherapy (or control IgG), measured the tumor volumes over time, and collected tumors at endpoint (Day 29) for immunophenotyping analysis. We found that while loss of Acod1 alone did not alter the growth of PRp tumors *in vivo,* it did restore sensitivity of these tumors to αPD-1 ICB therapy (Fig. 3c). To gain insight into how tumor growth was restricted in these tumors generated from Acod1-deficient cells, we characterized the immune infiltration in the tumors at the time of harvest (see schematic in Fig. S4a). While Acod1 loss was not sufficient to cause an increase in the total immune infiltrate within the tumor, the αPD-1 treatment, as expected, did augment the total abundance of immune cells (Fig. S4b). Moreover, neither knockout of Acod1 nor administration of αPD-1 monotherapy altered the intratumoral percentage of CD4^+^ T cells (Fig. 3d, e). However, the tumors generated from PRp cells deficient in Acod1 had a marked increase in the number of CD8^+^ T cells (Fig. 3d, f). Tumor regression following therapeutic αPD-1 is dependent upon pre-existing CD8^+^ T cells within the tumor underlying the importance of increased CD8^+^ T cell infiltrate in the Acod1-deficient tumors^45^. We therefore conclude that the reduction of Acod1 in cancer cells increases sensitivity to αPD-1 monotherapy and elevates cytotoxic CD8^+^ T cell numbers.

**Figure 3:**
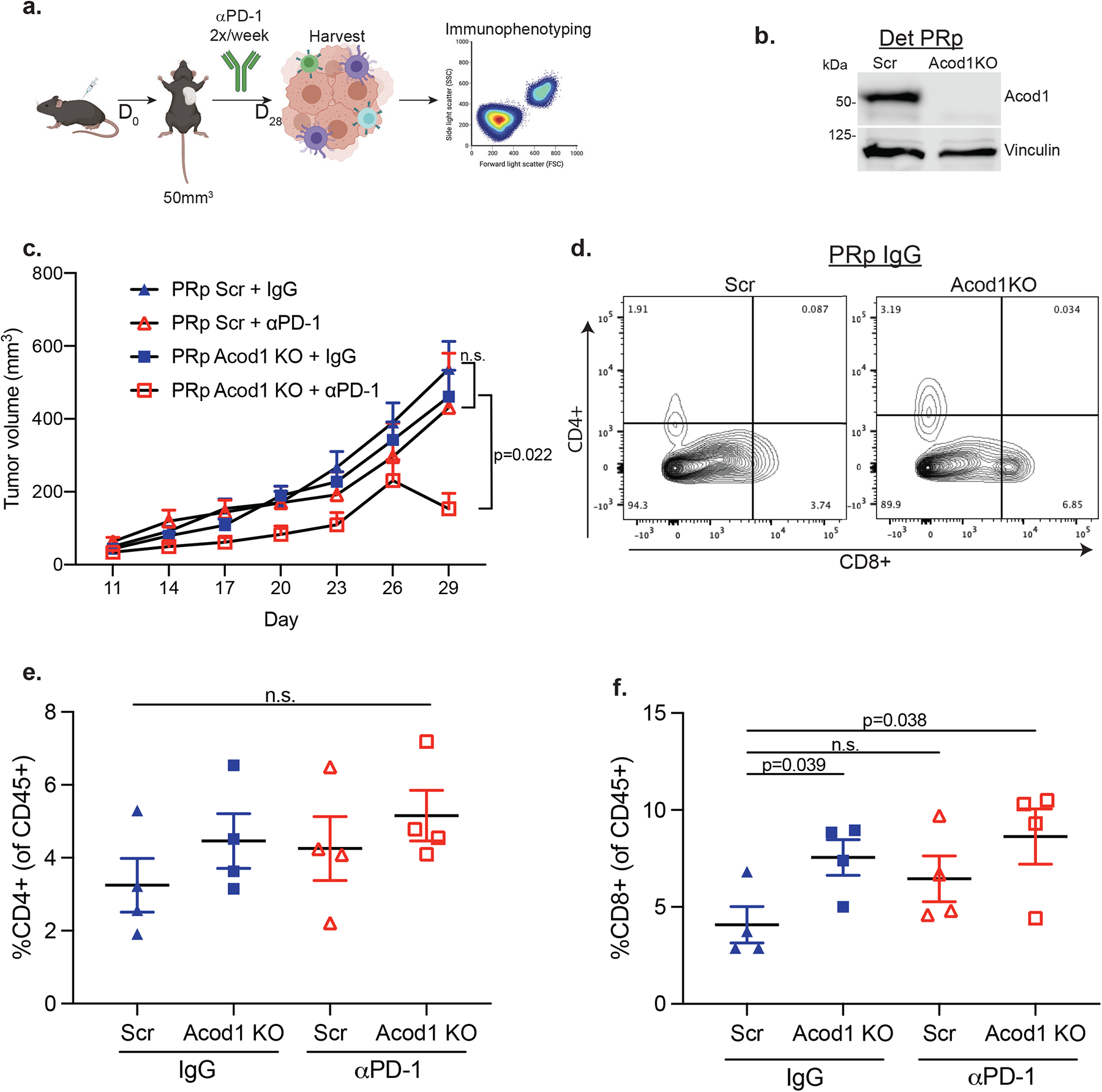
ACOD1 dictates sensitivity to αPD-1 ICB *in vivo*. **a**. Experimental design of *in vivo* experiment. **b**. Western blot for Acod1 of cells injected for tumor experiments. **c**. PRp-Scr or PRp-Acod1KO cells were subcutaneously injected in mice and received either isotype control (IgG) or αPD-1 mAb when tumors reached ≥50 mm^3^. Data are mean ± SEM (*n* = 6 to 10 per group) and *P* values were calculated by two-way ANOVA. **d**. Flow cytometry plots for CD8 versus CD4 from PRp-Scr (left) or PRp Acod1KO (right) tumors. **e**,**f**. Percentage of CD4^+^ (**e**) or CD8^+^ (**f**) T cells within each tumor. Data represent the mean ± SEM (*n* = 4 mice/group) and *P* values were calculated by two-tailed Student’s *t* test. n.s., represents ‘not significant’.

### Conditioned media from cancer cells with high levels of Acod1 inhibits naïve CD8^+^ T cell activation and proliferation

Given the increased number of CD8^+^ T cells in tumors derived from cells engineered to be deficient in Acod1, we reasoned that Acod1 may facilitate the release of immunomodulatory factors from cancer cells. More specifically, cancer cells that form αPD-1-resistant tumors have been shown to release immunosuppressive factors that can inhibit the transition of CD8^+^ T cells from a naïve to an activated state^46^, which could alter the sensitivity of tumors to αPD-1 ICB monotherapy. We therefore employed a T cell activation assay to investigate how PRp cell-conditioned media (CM) impacted CD8^+^ T cell activation and proliferation. Indeed, αCD3/CD28-stimulated CD8^+^ cells activated in CM generated from PRp cells had substantially reduced proliferation, as evidenced by both reduced EdU incorporation (Fig. 4a) and violet proliferation dye 450 (VPD450) dilution (Fig. 4b), whereas there was no deficit in proliferation of T cells activated in CM generated from PRs cells (Fig. 4a, b). In addition, T cell activation, as measured by cell surface levels of CD44 and CD25, was also dampened when naïve CD8^+^ T cells were activated in PRp (but not PRs) CM (Fig. 4c, d). Given that Acod1 levels are high in PRp cells and that reducing Acod1 sensitized PRp tumors to αPD-1 monotherapy, we posited that Acod1 may regulate the ability of PRp cells to block the activation and proliferation of CD8^+^ T cells. Additionally, previous work has demonstrated that PC3 cells (which have detectable ACOD1 levels), but not DU-145 cells (which do not have detectable ACOD1) (Fig. S3a), can secrete factors that modulate the activity of T cells^46^. We first confirmed that, as expected, our PRp-Acod1KO cells lost the capacity to both produce and secrete ITA (Fig. S5a, b). To interrogate the contribution of Acod1 to the inhibitory phenotype, CM was generated from these cell lines and used in the T cell proliferation/activation assays described above. Interestingly, CM generated from PRp-Acod1KO cells was unable to inhibit CD8^+^ T cell proliferation (Fig. 4e, 4f). Loss of Acod1 also increased activation markers (CD44/CD25) in T cells cultured in PRp-Acod1KO CM compared to PRp-Scr CM (Fig. 4g, h). Taken together, these data suggest that Acod1 regulates the secretion of inhibitory factors from cancer cells that negatively impact both activation and proliferation of CD8^+^ T cells.

**Figure 4:**
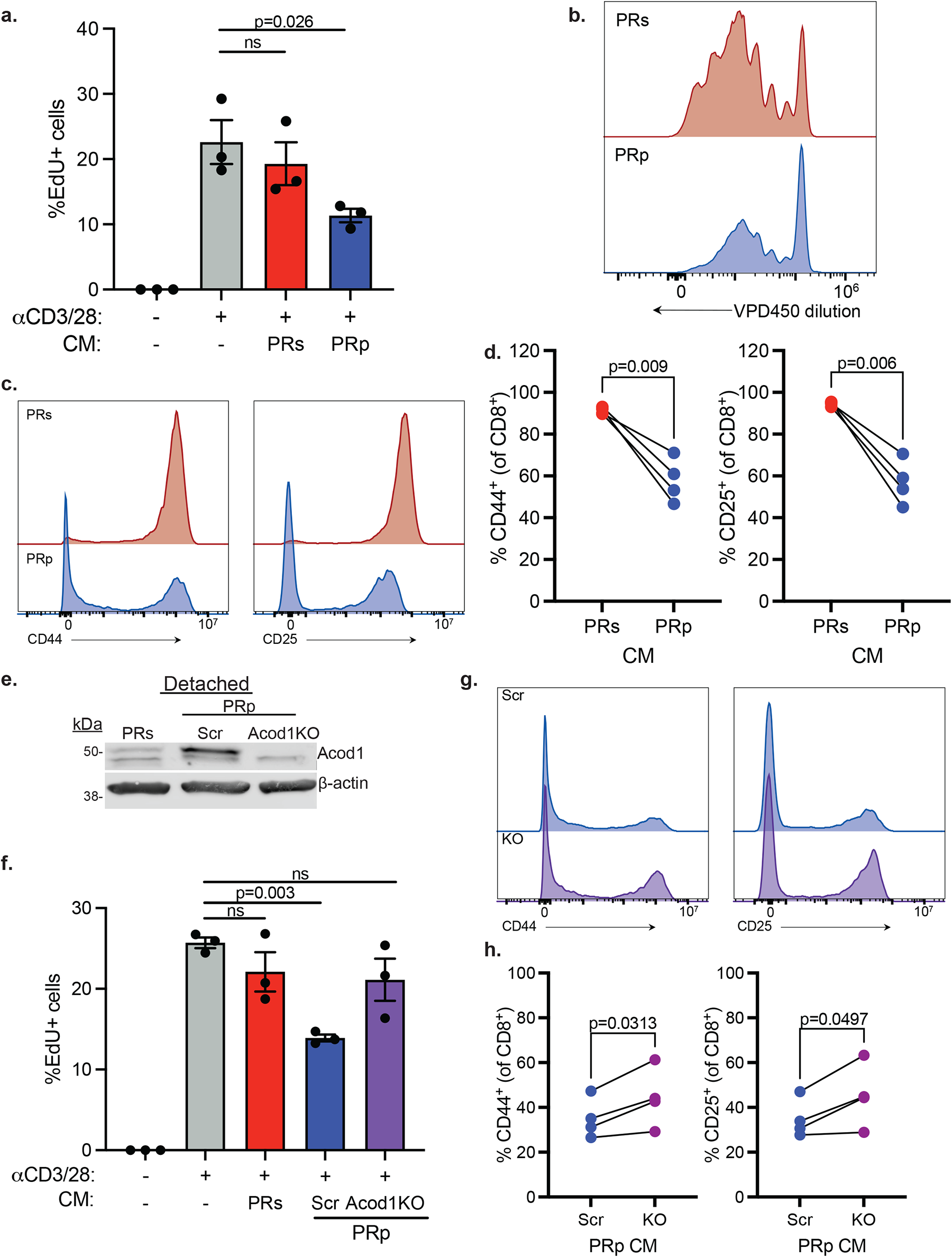
Secreted factors from ECM-Det ACOD1 high cells restrict T cell activation and proliferation. **a**. Percentage of EdU^+^ CD8^+^ T cells following 48 h of activation with αCD3/CD28 in indicated conditioned medium (CM). **b**. Representative histograms of violet proliferation dye 450 (VPD450) dilution in CD8^+^ T cells following 72 h of activation with αCD3/CD28 in indicated CM. **c**. Histograms of CD44 (left) and CD25 (right) expression in CD8^+^ T cells following 72 h of activation with αCD3/CD28 in indicated CM (*n* = 3). **d**. Percentage of CD44^+^ (left) and CD25^+^ (right) CD8^+^ T cells following 72 h of activation with αCD3/CD28 in indicated CM. **e**. Cells grown for 24 h in Det conditions. Lysates were collected and immunoblotted as noted. **f**. Percentage of EdU^+^ CD8^+^ T cells as described in (**a**). **g**. Histograms of CD44 (left) and CD25 (right) expression as described in (**c**). Scr = PRp-Scr CM; KO = PRp-Acod1 KO CM (*n* = 3). **h**. Percentage of CD44^+^ (left) and CD25^+^ (right) CD8^+^ T cells as described in (**d**). Data from EdU experiments (**a**, **f**) represent the means ± SEM of triplicate wells and *P* values were calculated by one-way ANOVA analysis. Data are representative of three independent experiments. The *P* values for the T cell activation experiments (**d**, **h**) were calculated using a paired, two-tailed *t* test (*n* = 4 mice/group). n.s., represents ‘not significant’. Western blotting and VPD450 dilution experiments were independently repeated a minimum of three times with similar results.

### Acod1-mediated inhibition of CD8^+^ T cell proliferation/activation is independent of extracellular ITA

As previously mentioned, the only documented role (to our knowledge) for Acod1 is catalyzing the decarboxylation of *cis*-aconitate to form ITA. Acod1-mediated ITA can be secreted into the extracellular space^18, 47^, and, despite its hydrophobic nature, can cross the plasma membrane and be utilized by recipient cells^48^. In addition to the known immunomodulatory roles of ITA in macrophages^44^, recent studies have examined a role for ITA in the regulation of T cell proliferation and differentiation^29, 30, 49^. We thus posited that secretion of extracellular ITA by cancer cells expressing Acod1 can result in the inhibition of CD8^+^ T cells proliferation/activation. To test this hypothesis, we first activated CD8^+^ T cells in media supplemented with increasing concentrations of exogenous ITA. Our choice of concentrations to test was informed by previous research assessing the abundance of extracellular ITA in different settings^47^. Surprisingly, exogenous ITA supplementation did not impact the proliferation of CD8^+^ T cells except at very high (10 mM) concentrations (Fig. 5a). Additionally, there was no impact of ITA on the percentage of activated CD8^+^ T cells at any of the concentrations tested (Fig. 5b,c); however a slight decrease in CD25 (but not CD44) on a per cell basis was noted with increasing ITA concentrations. In addition to examining activation and proliferation of CD8^+^ T cells, we also measured the effect of exogenous ITA supplementation on cytokine production. We observed a concentration-dependent decrease in interferon gamma (IFN-γ) production with increasing concentrations of ITA (Fig. 5d), but observed little to no effect on Granzyme B production (Fig. S6). As such, our data show that ITA does not phenocopy the effects of PRp CM on CD8^+^ T cell proliferation or activation, but impairs IFN-γ production by CD8^+^ T cells. Lastly, we thought it prudent to quantitatively determine the concentration of ITA in the CM generated from the PRs and PRp cell lines. As expected, we did see a significant elevation in the abundance of ITA secreted by PRp (compared to PRs) cells (Fig. 5e). However, the concentration of ITA in PRp CM topped out at slightly over 500 nM, far below the concentrations (10 mM) at which we observed effects of ITA on CD8^+^ T cell proliferation. Taken together, our data suggest that secretion of ITA is not responsible for the capacity of CM collected from Acod1-expressing cancer cells to antagonize CD8^+^ T cell proliferation and activation.

**Figure 5:**
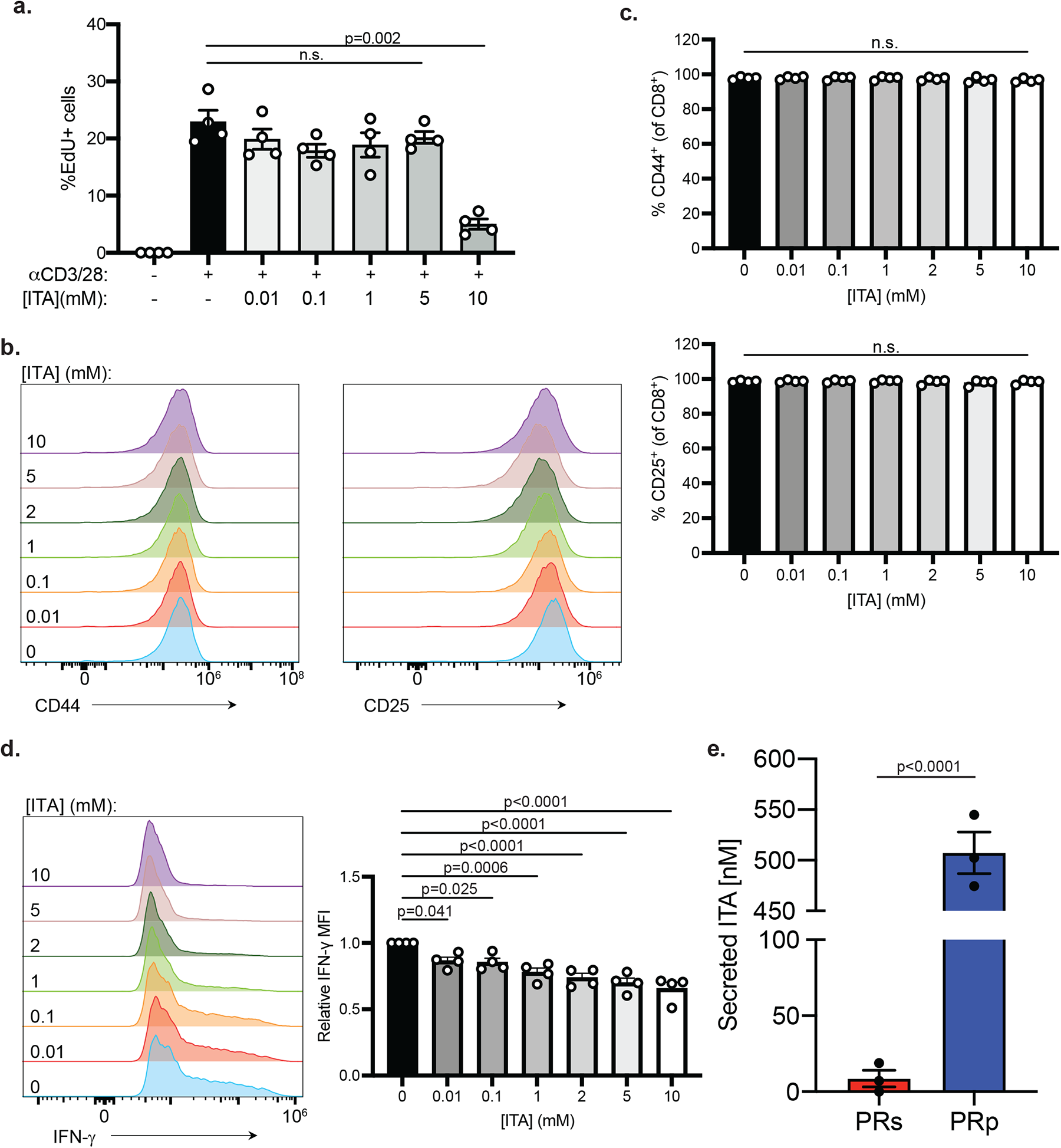
ACOD1-mediated effect on CD8^+^ T cells is independent of extracellular itaconate. **a**. Percentage of EdU^+^ CD8^+^ T cells following 48 h of activation with αCD3/CD28 in the presence of the indicated concentrations of itaconate (ITA). **b**. Histograms of CD44 (left) and CD25 (right) expression in CD8^+^ T cells following 72 h of activation with αCD3/CD28 in the presence of the indicated concentrations of ITA. **c**. Bar graphs showing the percentage of CD44^+^ (top) or CD25^+^ (bottom) CD8^+^ T cells activated as described in (**b**). **d**. IFN-γ production by CD8^+^ T cells activated as in (**b**). Representative histograms of IFN-γ expression in CD8^+^ T cells (left) and relative MFI of IFN-γ in IFN-γ^+^ CD8^+^ T cells (right). **e**. Concentration of ITA in the CM from PRs or PRp cells. Data in EdU experiment (**a**) represent the means ± SEM of four replicate wells. Data are representative of three independent experiments. Data in (**c-d**) represent the means ± SEM (*n* = 4 mice/group). one-way ANOVA analysis. Data in (**e**) are analyzed by two-tailed Student’s *t* test. n.s., represents ‘not significant’.

### Acod1 regulates the secretion of immunomodulatory small peptides

Our data demonstrate that cancer cells are capable of secreting factors that inhibit CD8^+^ T cells through a mechanism dependent on Acod1, but independent of extracellular ITA (see Fig. 4, Fig. 5). As such, we were interested in determining the identity of the secreted factor (or factors) that impacts CD8^+^ T cells. To do so, we first boiled CM from PRs or PRp cells to assess whether something proteinaceous was involved in the inhibition of CD8^+^ T cells. Interestingly, the capacity of CM from PRp cells to inhibit the proliferation of CD8^+^ T cells was maintained after boiling (Fig. 6a, S7a), suggesting that heat-labile proteinaceous factors were not responsible for the inhibitory effect. Next, we filtered CM from PRp cells to remove any factors that are larger than 3 kDa. The CM depleted of factors larger than 3 kDa retained its ability to block proliferation of CD8^+^ T cells (Fig. 6b). To rule out the possibility of protein fragments in the CM playing a role in inhibition of T cells, we treated PRp CM (that had been depleted of factors larger than 3 kDa) with Proteinase K to digest any remaining protein fragments. Surprisingly, Proteinase K addition was sufficient to rescue proliferation of CD8^+^ T cells activated in PRp CM (Figure 6c). Motivated by these data, we reasoned that small peptides, which can be secreted by cells and have been shown to exert a variety of biological functions^50–53^, may impact CD8^+^ T cell proliferation. In support of this possibility, treatment of PRp CM with dextran-coated charcoal (DCC), which can bind to and remove a diversity of small molecules, rescued the proliferation of CD8^+^ T cells exposed to PRp CM (Fig. 6d).

**Figure 6:**
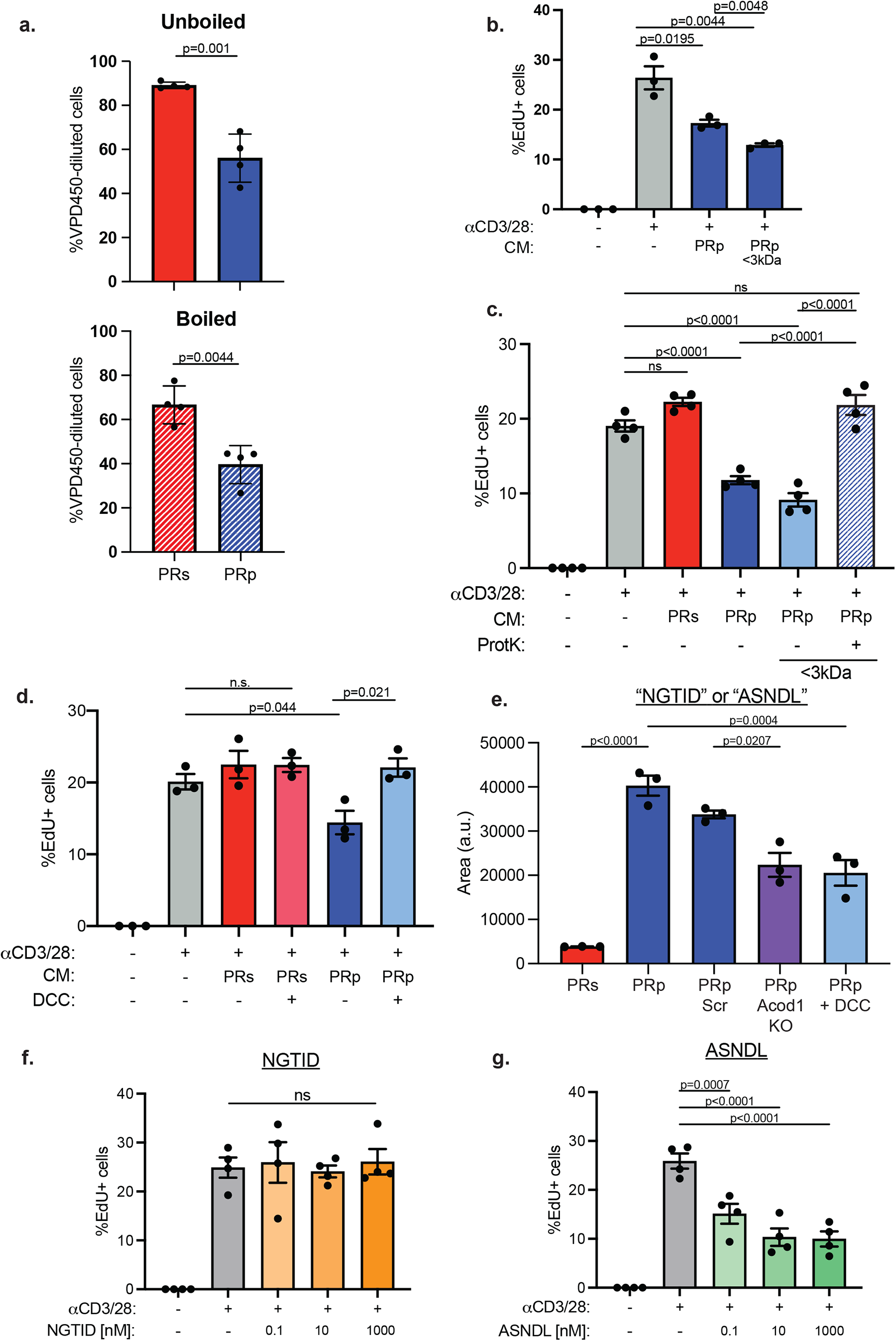
ACOD1 regulates the secretion of immunomodulatory peptides. **a**. Violet proliferation dye 450 (VPD450) dilution in CD8^+^ T cells following 72 h of activation with αCD3/C28 in indicated conditioned medium (CM) that was either unboiled (top) or boiled (bottom) prior to activating the T cells. **b**. Percentage of EdU^+^ CD8^+^ T cells following 48 h of activation with αCD3/C28 in full PRp CM or <3 kDa fraction of PRp CM. **c,d.** Percentage of EdU^+^ CD8^+^ T cells in indicated conditions following activation as noted in (**b**). **e**. Bar graph showing the relative abundance of a 518 Da molecular weight compound in the listed CM. **f,g**. Percentage of EdU^+^ CD8^+^ T cells following 48 h of activation in the presence of either NGTID peptide (**f**) or ASNDL peptide (**g**). Peptides used at 0.1, 10, or 1000 nM. Data in (**a**) represent the mean ± SEM (*n* = 4 mice/group). Data in EdU experiments (**b-d, f-g**) represent the means ± SEM of four replicate wells. Data are representative of three independent experiments. *P* values were calculated by two-tailed Student’s *t* tests (**a**) or one-way ANOVA analysis (**b-g**). n.s., represents ‘not significant’.

Given the likelihood of a small peptide in modulating CD8^+^ T cell proliferation, we sought to identify the peptide (or peptides) secreted by cells expressing Acod1. We therefore conducted mass spectrometric analysis on CM collected from different conditions. Since the factor would be more abundant in inhibitory CM, we sorted our results for hits where the relative abundance of the compound in PRp CM was at least 2-fold higher than PRs. We then looked for candidates that were higher in the inhibitory CM (PRp, PRp-Scr) and lower in the permissive conditions (PRs, PRp-Acod1KO, PRp+DCC) (Fig. S7b). Interestingly, we discovered a candidate (m/z = 519.2411) that was elevated in PRp CM compared to PRs CM which was also decreased in PRp CM when Acod1 is lost or when the CM is treated with DCC (Fig. 6e). Chemical formula prediction (Tracefinder v 1.8.63.0, Thermo Scientific) from accurate mass MS1 data considering C, H, O, N, and S atoms and isotopologue pattern matching revealed nine chemical formulae possibilities within +/-2 ppm MS1 mass error (Figure S7c). Since we hypothesized that this unknown is a peptide, we further ruled out eight of these matches that contained at least one S since there is no possible peptide sequences containing methionine or cysteine, the only amino acids with sulfur, that have the mass of interest. This remaining putative chemical formula was C_20_H_35_O_10_N_6_. Orthogonally, we searched the unknown m/z of 519.5411 using the METLIN detabase (excluding compounds with chlorine and/or fluorine atoms), an MS/MS experimental database over 700,000 molecular standards^54^. This revealed two possible compounds, both with the same chemical formula of C_20_H_35_O_10_N_6_ as was revealed by chemical formula prediction, which are pentapeptides NGTID or ASNDL (we were not able to resolve the order of the amino acids in these peptide sequences). Since spectral libraries do not exist for pentapeptides, preventing MS2 spectral matching, we opted to use a functional validation of this putative compound identification. We subsequently synthesized each of these pentapeptide species to assess their capacity to impact CD8^+^ T cell proliferation. While NGTID treatment did not have a discernible impact on CD8^+^ T cell proliferation (Fig. 6f), treatment with ASNDL led to a concentration-dependent reduction in CD8^+^ T cell proliferation (Fig. 6g). Taken together, our data suggest that Acod1 expression in cancer cells can lead to the secretion of immunomodulatory pentapeptide species that can negatively regulate proliferation in CD8^+^ T cells.

## Discussion

We discovered that cancer cells resistant to αPD-1 ICB have profound changes in mitochondrial metabolism that impact the immune microenvironment. More specifically, αPD-1 resistant cancer cells were discovered to have high levels of ITA owing to a significant elevation in Acod1. Engineering αPD-1 resistant cancer cells to be deficient in Acod1 restores the sensitivity of tumors arising from these cells to αPD-1 ICB monotherapy. Mechanistically, we found that αPD-1 resistant cancer cells secrete factors into the extracellular environment that negatively impact both activation and proliferation of CD8^+^ T cells. Surprisingly, this effect on CD8^+^ T cells is not due to the secretion of ITA. Instead, αPD-1 resistant cancer cells secrete small peptides that antagonize CD8^+^ T cell proliferation.

Our data reveal a new and important role for Acod1 in cancer cells and in the regulation of anti-tumor immunity. As mentioned previously, *Acod1* expression has largely been studied in the myeloid lineage (e.g. monocytes, macrophages). However, our findings add to a growing body of evidence suggesting that the activity of Acod1 can alter the TME in a fashion that alters ICB sensitivity^44^. In particular, we report here that Acod1 is expressed in both human and mouse cancerous epithelial cells. Similarly, our data add to the growing body of literature focused on understanding the immunomodulatory properties of ITA, including recent work that links ITA to the modulation of regulatory T cell (Treg) differentiation^29^ and to the regulation of CD8^+^ T cells^30^. The latter study presents intriguing data suggesting that ITA secreted by myeloid-derived suppressor cells (MDSCs) can be taken up by CD8^+^ T cells to negatively impact activation and proliferation. While we did not observe evidence of ITA-mediated changes in CD8^+^ T cell activation or proliferation (see Figure 5), we focused our analysis on a much lower concentration that was based on our ITA measurements in CM from PRp cells (in the nM range). It is thus possible that MDSC-derived ITA exists in higher concentrations in the extracellular environment and could be somehow distinct from cancer cell-derived ITA. Relatedly, immunomodulatory derivatives of ITA (e.g. citraconate and mesaconate) have recently been discovered^55, 56^. While the mechanisms by which these derivatives are generated remain elusive, it is possible that there are important differences in ITA derivatization in distinct cell types that may impact its biological activities.

Our data suggest that the activity of Acod1 can cause cancer cells to secrete small, immunomodulatory peptides that negative impact CD8^+^ T cell proliferation. More specifically, we discovered the pentapeptide ASNDL in CM collected from cancer cells with elevated Acod1. While our data demonstrate that ASNDL production is dependent on Acod1 levels in cancer cells and that it can inhibit CD8^+^ T cell proliferation, it remains possible that there are other small peptides with similar effects on CD8^+^ T cells. Future studies aimed at systematically studying the configuration of immunomodulatory small peptides could be informative in leveraging these findings for clinical translation. Additionally, the mechanism by which Acod1 leads to the generation and secretion of peptides like ASNDL will require future study. Intriguingly, a recent study discovered that Acod1-mediated ITA can promote lysosomal biogenesis through a mechanism dependent on transcription factor EB (TFEB)^57^. It thus seems plausible that ITA-mediated changes in lysosomal activity could impact protein degradation in a fashion that produces immunomodulatory peptides like ASNDL.

## Acknowledgements

We thank Veronica Schafer and all current/past Schafer lab members for helpful comments and/or valuable discussion. We thank Jianneng Li (Notre Dame) for the DU-145 cells, Sara Cole (Notre Dame Integrated Imaging Facility) for assistance with confocal microscopy, and the staff at the Freimann Life Science Center for assistance with animal care. ZTS is supported by the National Institutes of Health/National Cancer Institute (R01CA262439), the Coleman Foundation, the Malanga Family Excellence Fund for Cancer Research at Notre Dame, the College of Science at Notre Dame, the Department of Biological Sciences at Notre Dame, and funds from Mr. Nick L. Petroni. XL (R01CA248033, R01CA280097) and SM (5F99CA274694) are also supported by the National Institutes of Health/National Cancer Institute. RGJ is supported by the Paul G. Allen Frontiers Group Distinguished Investigator Program, National Institutes of Health/National Institute of Allergy and Infectious Diseases (R01AI165722), and VAI.

## Materials and Methods

### Cell Culture

MDA-MB-436, PC3, and GCT were purchased from ATCC. DU145 cells were a kind gift from Jianneng Li (Notre Dame). PRs and PRp were developed from a spontaneous prostate tumor of *PB-Cre^+^ Pten^L/L^ Tp53^L/L^ Smad4^L/L^* transgenic mice^32^. MDA-MB-436, PRs, and PRp were cultured in DMEM (Gibco). DU145 and PC3 were cultured in RPMI-1640 (Gibco). GCT cells were cultured in McCoy’s 5A Medium (Gibco). All media were supplemented with 10% FBS (Invitrogen) and cells were cultured in the presence of 1% penicillin/streptomycin (HyClone). All cell lines were cultured in a humidified incubator at 37^°^C with 5% CO_2_. All cells were tested routinely for mycoplasma-free status by PCR kit (Bulldog Bio, 25234).

### Lentiviral transduction

Lentiviral constructs with gRNA targeting mouse Acod1 (5’-GGGCTTCCGATAGAGCTGTG-3’ and 5’-AACGTTGGTATTGAAGTACA-3’), or SCR gRNA (5’-GTGTAGTTCGACCATTCGTG-3’) in the puromycin-selectable pLV-hCas9 vector were purchased from VectorBuilder (Chicago, IL). HEK293T cells were transfected with target DNA along with the packaging vectors psPAX2 and pCMV-VSV-G using Lipofectamine 2000 (Invitrogen). Virus was collected 48h after transfection, filtered through a 0.45 μm filter (EMD Millipore), and used for transduction of PRp cells in the presence of 8 μg ml^-1^ polybrene. Stable polyclonal populations of cells were selected using 3 μg ml^-1^ puromycin (Invitrogen).

### Mitochondrial measurements

Mitochondrial abundance was measured with MitoTracker Red CM-H_2_XRos (Invitrogen, M7513). Cells were washed with PBS and stained with 100nM of MitoTracker in growth media for 30 mins at 37^°^C. After incubation, cells were either counterstained with Hoechst33342 (Invitrogen, H1399) and fixed with 4% PFA for 15 mins for analysis by confocal fluorescence microscopy or washed with PBS and analyzed by flow cytometry. Confocal microscopy was conducted using a Nikon A1R-MP microscope (Nikon, Melville, NY). Maintenance of mitochondrial pool was assessed by the median fluorescence intensity (MFI) ratio of each cell line using the attached conditions as an internal control.

### Immunoblotting

Attached and ECM-detached cells were washed once with ice cold PBS and lysed in RIPA buffer (50 mM Tris (pH 8.0), 150 mM NaCl, 1% Nonidet P-40, 0.5% Sodium deoxycholate, 0.1% SDS) supplemented with 1 mg ml^-1^ aprotinin, 5 mg ml^-1^ leupeptin, 20 mg ml^-1^ phenylmethylsulfonyl fluoride (PMSF) and HALT phosphatase inhibitor mixture (Thermo Fisher Scientific). Lysates were collected after centrifugation for 15 min at 4°C at 14,000 r.p.m. and normalized by BCA assay (Pierce Biotechnology). Normalized lysates underwent SDS-PAGE and transfer/blotting were carried out as described previously^58^.

The following primary antibodies were used for western blotting: β-actin (Sigma-Aldrich; no. A1978) (1:10000), Vinculin (Proteintech; 66305-1-Ig) (1:3000), mouse Irg1/Acod1 (Cell Signaling Technology (CST); no. 17805) (1:1000), human IRG1/ACOD1 (CST; no. 77510) (1:1000), Tim23 (BD Biosciences; no. 611222 (1:2000), Tom20 (CST; no. 42406) (1:1000). Secondary antibodies used were Alexa Fluor Plus 680 and 800 (Thermo Fisher Scientific, no. A32788, A32808) (1:10000) against mouse and rabbit, respectively, and bands were visualized with the LiCor Odyssey CLx (Licor).

### RNA isolation and quantitative real-time PCR

Total RNA was isolated with Zymo Quick RNA Miniprep kit (Zymo Research, R1054). RNA (1μg) was reverse transcribed into complementary DNA using an iScript Reverse Transcription Supermix kit (Bio-Rad, Hercules, CA, USA). The relative levels of gene transcripts compared with the control gene 18S were determined by quantitative real-time PCR using SYBR Green PCR Supermix (Bio-Rad) and specific primers on a 7,500 fast real-time PCR system (Applied Biosystems, Life Technologies, Waltham, MA, USA). Amplification was carried out at 95 ^°^C for 12 min, followed by 40 cycles of 15 s at 95 ^°^C, and 1 min at 60 ^°^C. Error bars represent s.e.m., and *P*-values were calculated using a one-way ANOVA. The fold change in gene expression was calculated as 2^−ΔΔCT^ and normalized to attached PRs cells. The primers used were as follows:

*Acod1:* F: GCAACATGATGCTCAAGTCTG and R: TGCTCCTCCGAATGATACCA 18S: F: GGCGCCCCCTCGATGCTCTTAG and R: GCTCGGGCCTGCTTTGAACACTCT

### Metabolic Tracing Experiments

Cells were cultured in their respective media in detached (6 mg ml^-1^ poly-HEMA coated) plates for 20 h. Cells were then collected, washed with PBS, and replated in either uniformly labeled ^13^C-Glucose (Cambridge Isotope Labs, CLM-1396) and unlabeled ^12^C-Glutamine (Sigma-Aldrich, 49419), or unlabeled ^12^C-Glucose (Sigma-Aldrich, G7021) and uniformly labeled ^13^C-Glutamine (Cambridge Isotope Labs, CLM-1822) and incubated in a 37^°^C humidified incubator with 5% CO_2_ for another 4 h. Cells were then centrifuged for 1 min at 900 RPM at 4^°^C to pellet the cells. Supernatant was collected and snap-frozen in liquid nitrogen. Cell pellets were washed briefly with ice-cold 0.9% NaCl, centrifuged to remove remaining media and snap-frozen in liquid nitrogen.

Metabolites were extracted with the Bligh-Dyer method^59^. The polar metabolite containing aqueous phase was dried and resuspended in 50 ml H_2_O. For glucose and glutamine tracing and profiling experiments, LC-MS analysis was conducted on an Orbitrap ID-X (Thermo) as previously described^60, 61^. Briefly, negative mode analytes were assessed using a Waters BEH Amide column (Mobile Phase A: 10mM ammonium acetate, 0.1% ammonium hydroxide, 0.1% medronic acid. Mobile phase B: 90% ACN, 10mM ammonium acetate, 0.1% ammonium hydroxide, 0.1% medronic acid). Positive mode analytters were separated with a Waters T3 column (Mobile phase A: water with 0.1% formic acid; Mobile Phase B: ACN with 0.1% formic acid). Data was analyzed using EL-Maven and Compound Discoverer (v 3.1, Thermo).

To confirm loss of ACOD1 activity in the Acod1KO cells, 4x10^5^ PRp-Scr or – Acod1KO cells/well were plated in 6-well poly-HEMA coated plates and stored in a 37 ^°^C humidified incubator with 5% CO_2_ for 24 h. After 24 h, cells were then collected, washed with PBS, and replated in uniformly labeled ^13^C-Glucose and placed back into the incubator for another 24 h. Cells and supernatant were then processed as previously stated and subjected to Bligh-Dyer metabolite extraction (above) to detect ^13^C-labeled-ITA in either the intracellular cell pellet or the secretome. Data were collected on an Orbitrap Exploris 240 mass spectrometer (Thermo) using a tributylamine ion-paired reversed phase chromatography as described previously^60, 61^. Data were analyzed in Compound Discoverer (v 3.3).

### Itaconate quantification

4x10^5^ PRs or PRp cells/well were plated in 6-well poly-HEMA coated plates in 4 ml of complete DMEM/well in triplicate and stored in a 37 ^°^C humidified incubator with 5% CO_2_ for 48 hours. Cells and supernatant were then processed as previously stated. A standard curve of neat itaconate (I29204, Sigma) was prepared using half-log dilutions from 316.23µM to 0.001µM, plus neat solvent as a zero blank. Equal volumes Samples and standards were extracted in 1mL of ice-cold acetonitrile:methanol:water (4:4:2 v/v). Data was collected using the tributylamine ion-paired reversed phase chromatography on the Orbitrap Exploris 240 a described above. Peak picking and integration was conducted in Skyline. Linear regression of standard curve peak areas were used to calculate ITA concentration in experimental samples.

### Untargeted metabolomics of conditioned media

To screen CM for peptide and other small-molecule candidate effectors, compounds were extracted from CM using a modified Bligh-Dyer extraction (200µL of conditioned media added to 690µL of 1:1 chloroform:methanol and 110 uL of water). The aqueous layer was collected, dried, and resuspended in 100µL of water. To maximize compound coverage, 4uL of each sample were injected twice on an Orbitrap ID-X, one for ESI positive mode and one for ESI negative mode, on each of the BEH amide and T3 chromatographies described above. Full scan data were analyzed in Compound Discoverer (v3.3) to identify candidate features. Candidate features were interrogated using chemical formula prediction (Tracefinder) and MS2 spectral matching (mzCloud, NIST2020).

### Chemical reagents

The NFκB inhibitor BAY1170-82 (Sigma-Aldrich, no. B5556) was used at 2.5 μM on ECM-detached PRp cells for 9 h in detachment. Cells were then collected and processed for immunoblotting or RNA extraction and qRT-PCR.

### Exogenous ITA experiments

Itaconic acid (ITA) (Sigma-Aldrich, no. I29204) was freshly made before use in the media used for the assay. The pH of the ITA solutions were adjusted to 7.4 using NaOH, and then used at the given concentrations listed in the experiment.

### Conditioned Media

Conditioned Media (CM) was generated by plating 4x10^5^ cells/well in a poly-HEMA coated 6-well plate in 4 ml total volume of respective media. After 48 h of incubation, we harvested the supernatant, centrifuged at 10,000g for 15 min at 24^°^C, collected the supernatant, aliquoted, snap-froze in liquid nitrogen, and stored at -80^°^C until ready to use in experiments. For all *in vitro* assays involving naïve CD8^+^ T cells, the CM was diluted 1:1 with fresh medium.

### T cell isolation

To isolate naïve CD8^+^ T cells, spleens from 6-8 week old C57BL/6N (Envigo) mice were harvested and single cell suspensions were generated by passing through a 70 μm filter (Falcon, 352350) then red blood cells were lysed with ACK lysis buffer (Gibco, A1049201) for 2 min on ice. Naïve CD8^+^ T cells were then purified from total lymphocytes by negative selection according to the manufacturer’s instructions (Miltenyi Biotec, 130-096-543) and cultured in DMEM containing 10% FBS, 1% penicillin/streptomycin, and 2-ME (Sigma-Aldrich). For *in vitro* analysis, naïve CD8^+^ T cells were stimulated with αCD3 and αCD28 mAb-coated dynabeads (Thermo Fisher Scientific, no. 11452D).

### EdU incorporation assay

EdU (5-ethynyl-2’-deoxyuridine) was diluted in media and added to cells for 4 h at 37^°^C in a humidified incubator with 5% CO_2_. After incubation, T cells were washed in 1x PBS, replated into a 96-well plate, and fixed using 4% PFA for 15 min. The Click-iT EdU kit (ThermoFisher, no. C10340) AF^+^647 was used according to the manufacturer’s instructions. Cells were counterstained with DAPI (1:2000) for 10 min and then imaged on a Zeiss inverted light microscope (Carl Zeiss, Axio Observer). Images were processed in ImageJ to get %EdU+, which was scored as (cells EdU+/cells DAPI+)*100.

### *In vitro* T cell activation assays

Naïve CD8^+^ T cells were purified from the spleens of female C57BL6/J mice (Jackson Laboratories) by negative selection (StemCell Technologies, 19858A). To evaluate cell proliferation by Violet Proliferation Dye 450 (VPD450) dilution, cells were stained with VPD450 according to the manufacturer’s instructions (BD Biosciences, 562158). 1.5x10^5^ naïve CD8^+^ T cells were seeded per well of a 96-well plate coated with αCD3ε (clone 145-2C11; 2 μg/mL) and αCD28 (clone 37.51; 1 μg ml^-1^) antibodies (eBioscience). Cells were seeded in 100 μL of Iscove’s Modified Dulbecco’s Medium (IMDM) supplemented with 10% Nu-Serum IV culture supplement (Corning), 1% penicillin/streptomycin, and 50 μM 2-ME. For assays involving CM, 2-ME was added to the CM at a final concentration of 50 μM, and then 100 μL of CM was added per well. For assays involving ITA, ITA was dissolved in IMDM and the pH adjusted to 7.4 prior to adding it to the cells at the indicated final concentrations. Cells were incubated at 37^°^C in a humidified incubator with 5% CO_2_ for 72 h. To assess cytokine production, cells were treated with phorbol 12-myristate 13-acetate (PMA; 50 ng ml^-1^) and ionomycin (500 ng ml^-1^) for 4 h at 37^°^C, with GolgiStop (BD Biosciences; 1:1500 dilution) added for the last 2 h of stimulation prior staining for flow cytometry. After PMA/ionomycin stimulation, the cells were stained with cell surface marker antibodies and a cell viability dye for 1 h at 4^°^C, fixed and permeabilized using the FoxP3/Transcription Factor Staining Buffer Set (eBioscience, 51-2092KZ) for 1 h at 4^°^C, and then stained with intracellular marker antibodies for 1 h at 4^°^C. The following antibodies were used for flow cytometry: CD8α-BUV395 (BD Biosciences, 563786), CD4-BV605 (BioLegend, 100548), CD25-Alexa Fluor 488 (eBioscience, 53-0251-82), CD44-BUV805 (BD Biosciences, 741921) IFN-γ-APC (eBioscience, 17-7311-82), Granzyme B-PE/Dazzle 594 (BioLegend, 372216). Cell viability was assessed by using Fixable Viability Dye eFluor 780 (eBioscience, 65-0865-14) according to the manufacturer’s protocol. Analytical flow cytometry was performed using a Cytek Aurora cytometer and data analysis performed using FlowJo (Tree Star) software.

### Heat-inactivation treatment of conditioned media

To heat-inactivate the polypeptide component of the supernatant, collected CM were boiled for 5 min at 100^°^C. After cooling, boiled CM was used 1:1 with fresh media for *in vitro* T cell assays as previously stated.

### Dextran-coated charcoal (DCC)

To remove small molecules, the CM were treated with DCC, which removes small molecules (e.g. nucleotides, lipids, peptides) by binding them on the surface. 12 mg of DCC per 500 μL of CM was added and incubated for 20 min at 25^°^C, followed by centrifugation at 10,000g for 30 min. After centrifugation, the supernatant that was cleared by the DCC was collected, snap-frozen in liquid nitrogen, and stored in -80^°^C until use.

### Fractionation of conditioned media and Proteinase K treatment

Culture supernatants were fractionated using Amicon ultra-centrifugal filters (Millipore Sigma, no. UFC900324) with 3 kDa cut-off. Supernatants were added to the 3 kDa filter and centrifuged at 12,000g for 30 min at 4^°^C. The filtered supernatant was collected and either stored as previously described or further treated with Proteinase K. For Proteinase K, 2 mg ml^-1^ of Proteinase K (Sigma-Aldrich, no. P2308) was added to filtered CM and incubated for 40 min at 37^°^C. Proteinase K-treated CM was added to a new 3 kDa filter and centrifuged at 12,000g for 30 min at 4 ^°^C to remove the Proteinase K.

### Mouse tumor experiment

All animal work performed in this study was approved (19-10-5588) by the Institutional Animal Care and Use Committee (IACUC) at the University of Notre Dame. Male C57BL/6N mice, 6 weeks old, were acquired from Envigo Laboratories. 5x10^5^ tumor cells were injected subcutaneously into the flank of 8 week-old mice. Once tumors reached ≥50 mm^3^, mice were i.p.-injected with 10 mg/kg of αPD-1 mAb (clone RMP1-14); the mAb treatment was repeated every third day. For control groups, an isotype control for the αPD-1 mAb (Rat IgG2a, κ) (clone RTK2758) was injected on the same schedule. Tumor sizes were measured every third day using a digimatic caliper (Louisware) and tumor volume was calculated using the formula [(length x width^2^)/2].

### Tumor immunophenotyping

Tumors were minced and digested in complete DMEM with 10% FBS and 1 mg/ml collagenase IV (Invitrogen, 17104019) at 37^°^C for 1 h, followed by passing through 40 μm strainers. Erythrocytes were lysed with ACK lysis buffer. Cells were treated with mouse Fc-block αCD16/32 (BioLegend, 101302) for 10 min, and stained with primary fluorophore-conjugated antibodies for 30 min. CD45-FITC (35-0451), CD4-APC (20-0041), and CD8-PE (50-1886) were used in the antibody cocktail (Tonbo Biosciences). Cells were25ashedm resuspended in Live/Dead Aqua 405 nm (Invitrogen) for 15 min, washed again, and then analyzed on a BD LSRFortessa X-20 cytometer.

### Peptide assays

NGTID and ASNDL peptides were synthesized to >95% purity by Peptide 2.0 (Chantilly, VA). Peptides stocks were resuspended in 50% MeOH and 50% ultra-pure H_2_O, aliquoted, and stored at -80 ^°^C. Each peptide was used fresh in the indicated assays and diluted to the working concentrations shown in their respective media. For most assays, peptides were used at 0.1, 10, and 1000 nM, unless otherwise indicated.

### Statistical Analysis

Statistical analyses were performed using GraphPad Prism v9.0. Unless otherwise mentioned, all data are presented as mean ± SEM (standard error of the mean). Sample sizes, error bars, *P* values, and statistical methods are denoted in figure legends. Statistical significance was defined as *P* < 0.05.

## Figure Legends

**Supplemental Figure S1:**
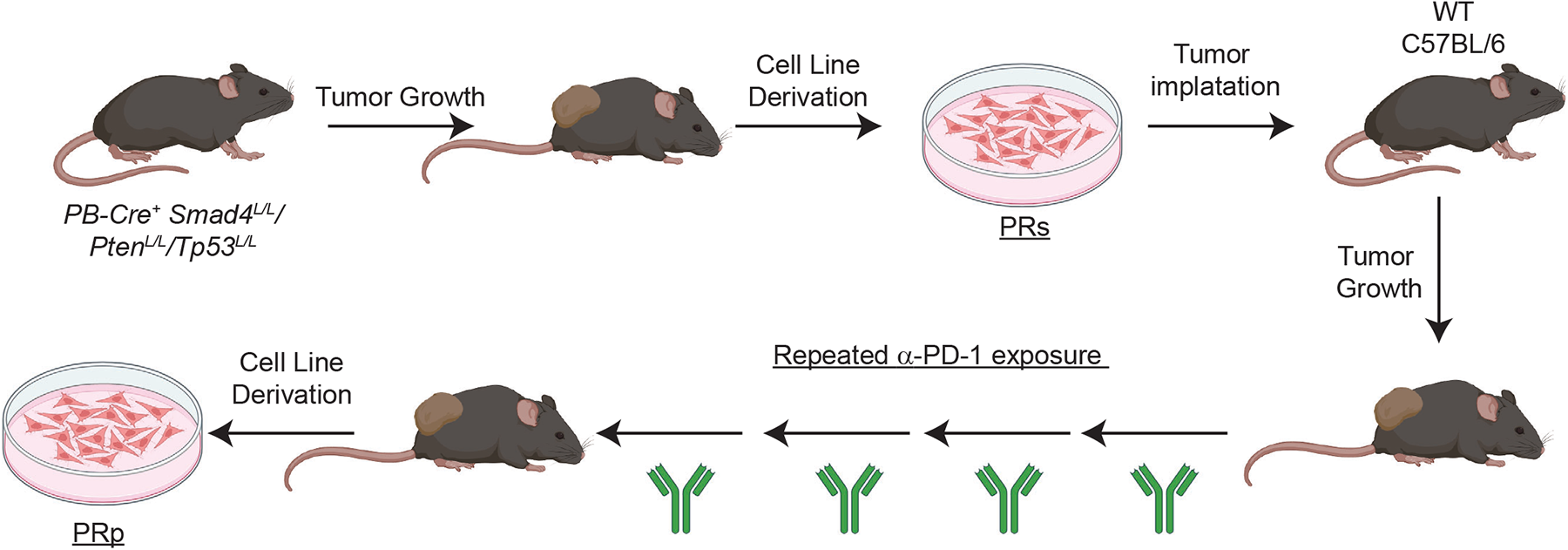
PRs and PRp cell line generation. Schematic of PRs and PRp cell line generation. PRs cells were established from a spontaneous prostate tumor. Serial exposure of residual PRs cells to αPD-1 mAb generated tumors that were completely unresponsive to αPD-1. PRp cell line was established from an αPD-1-unresponsive tumor.

**Supplemental Figure S2:**
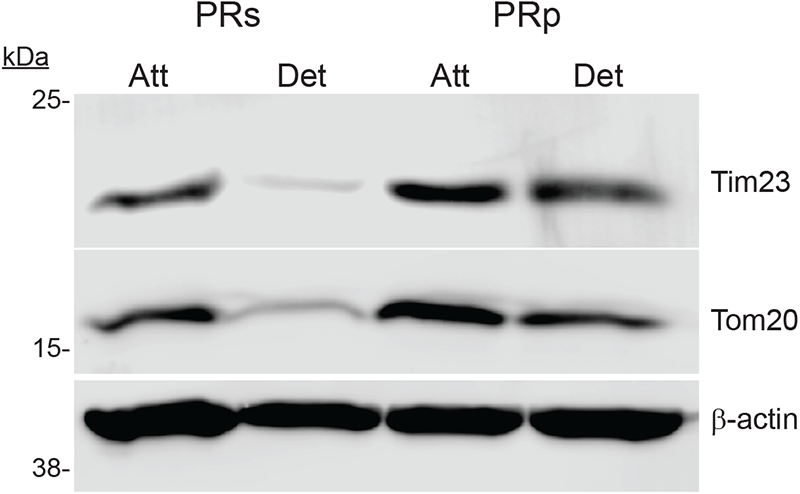
Mitochondrial protein depletion in ECM-Detachment. PRs and PRp cells were grown in the indicated conditions for 24 h. Lysates were collected and immunoblotted as noted. Western blotting experiments were independently repeated a minimum of three times with similar results.

**Supplemental Figure S3:**
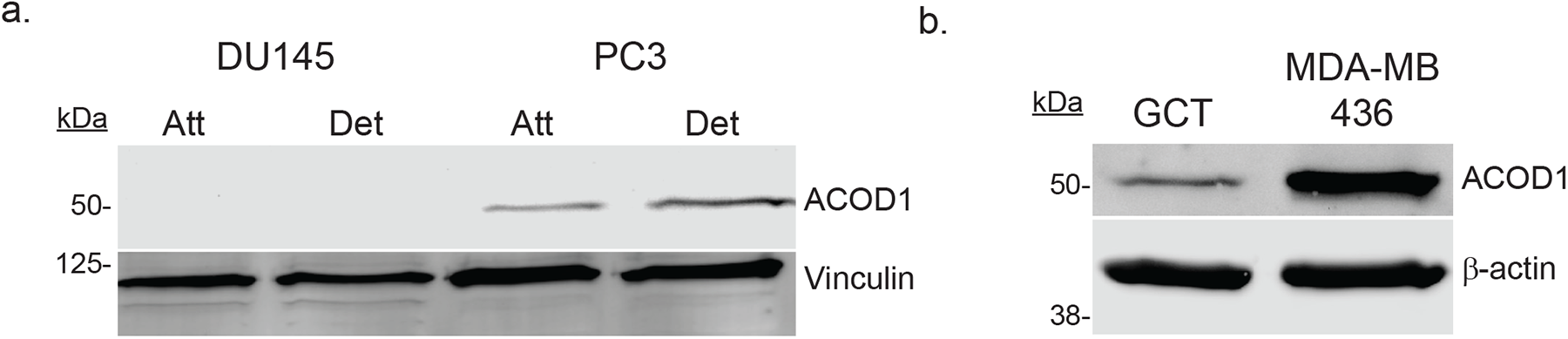
Human cancer cells have detectable ACOD1. **a**. DU145 and PC3 cells were grown in the indicated conditions for 24 h. Lysates were collected and immunoblotted as noted. **b**. GCT and MDA-MB-436 cells were grown in attached conditions for 24 h. Lysates were collected and immunoblotted as noted. Western blotting experiments were independently repeated a minimum of three times with similar results.

**Supplemental Figure S4:**
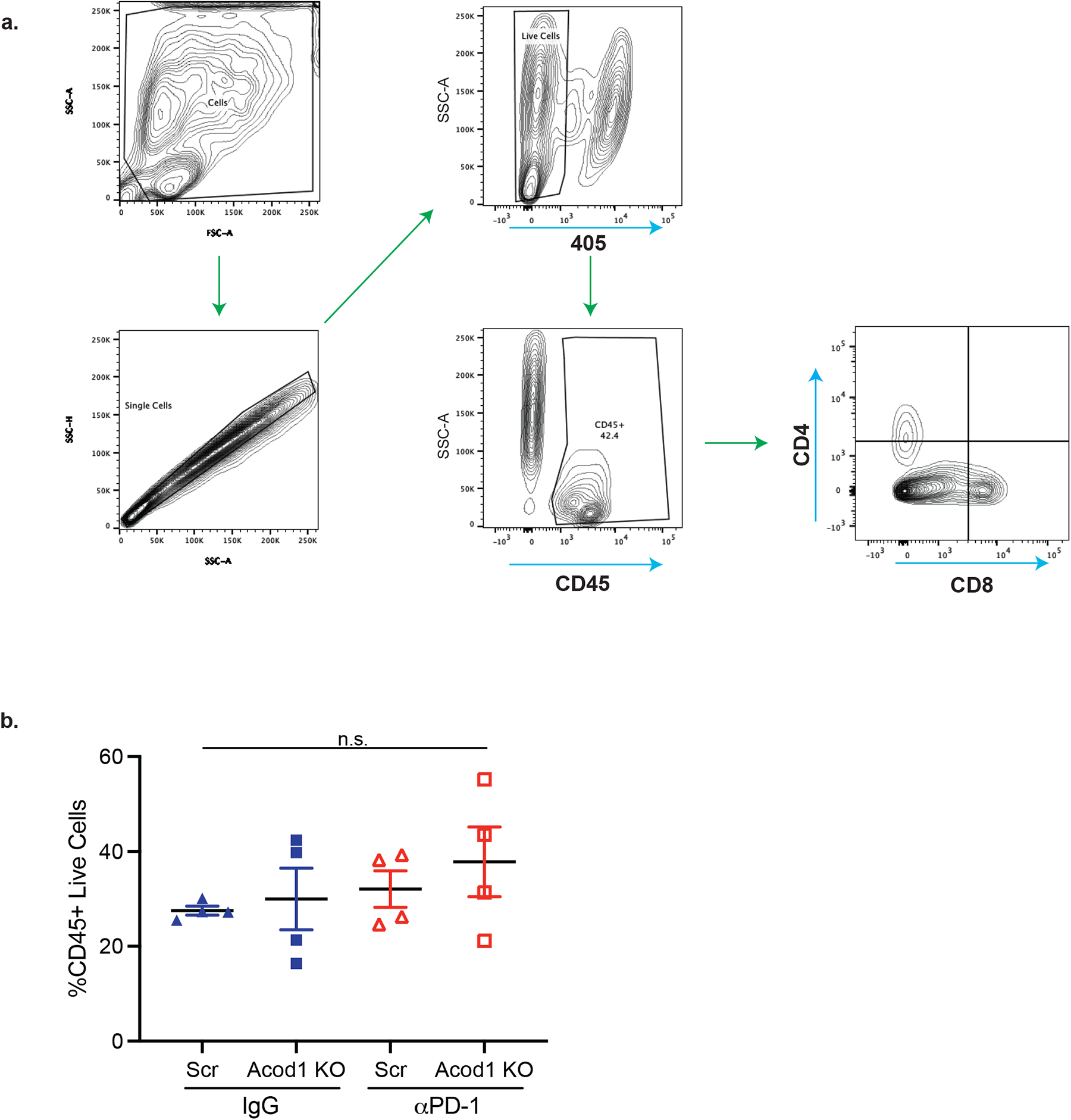
Flow plots for tumor experiment. **a**. Flow cytometry gating strategy for tumor-infiltrating lymphocytes (TILs). **b**. Percentage of immune cells (CD45^+^) from tumors.. Data are mean ± SEM (*n* = 4 mice/group) and *P* values were calculated by one-way ANOVA. n.s., represents ‘not significant’.

**Supplemental Figure S5:**
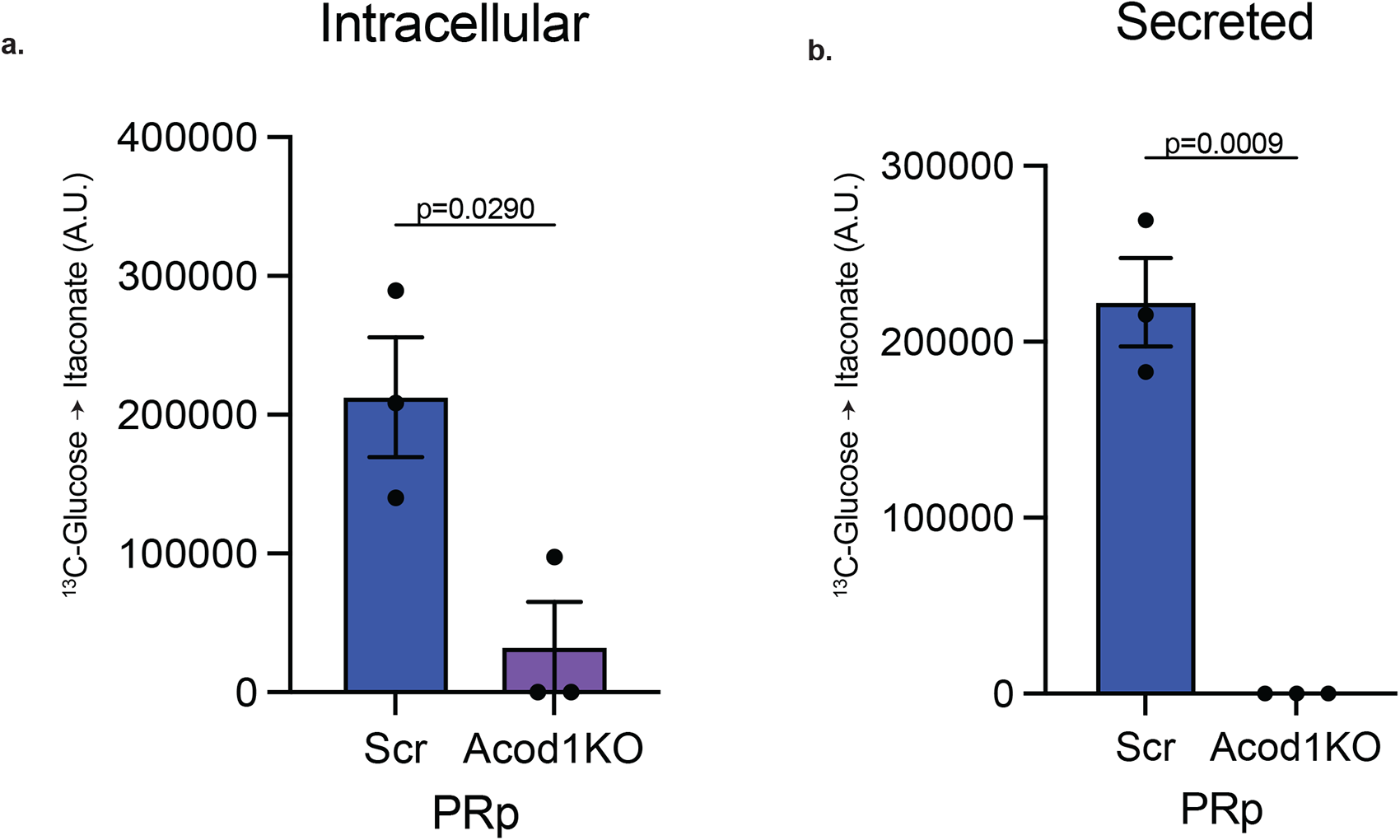
Acod1 KO reduces ITA production 24 h [U-^13^C]glucose tracing of PRp-Scr and PRp-Acod1 KO cells in ECM-Det conditions. **a**. Relative abundance of intracellular ^13^C-labeled itaconate (ITA). **b**. Relative abundance of extracellular/secreted ^13^C-labeled ITA found in the conditioned media. Graphs represent data collected from 3 biological replicates. *P* values are calculated by two-tailed Student’s *t* test. Data are mean ± SEM.

**Supplemental Figure S6:**
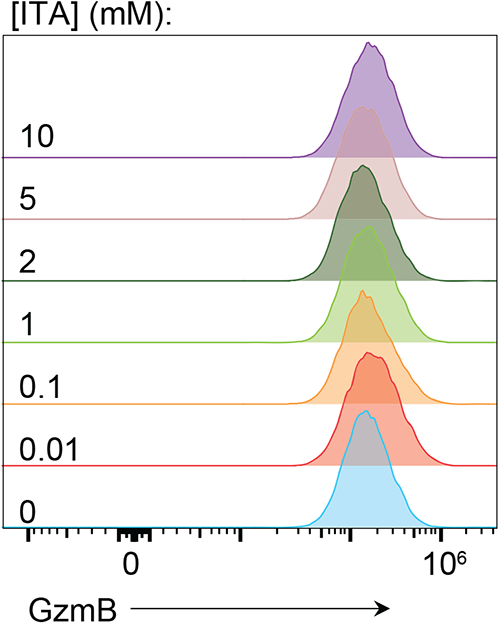
Granzyme B production by CD8^+^ T cells following activation in the presence of itaconate. Representative histograms of Granzyme B (GzmB) expression in CD8^+^ T cells following 72 h of activation with αCD3/CD28 in the presence of the indicated concentrations of ITA.

**Supplemental Figure S7:**
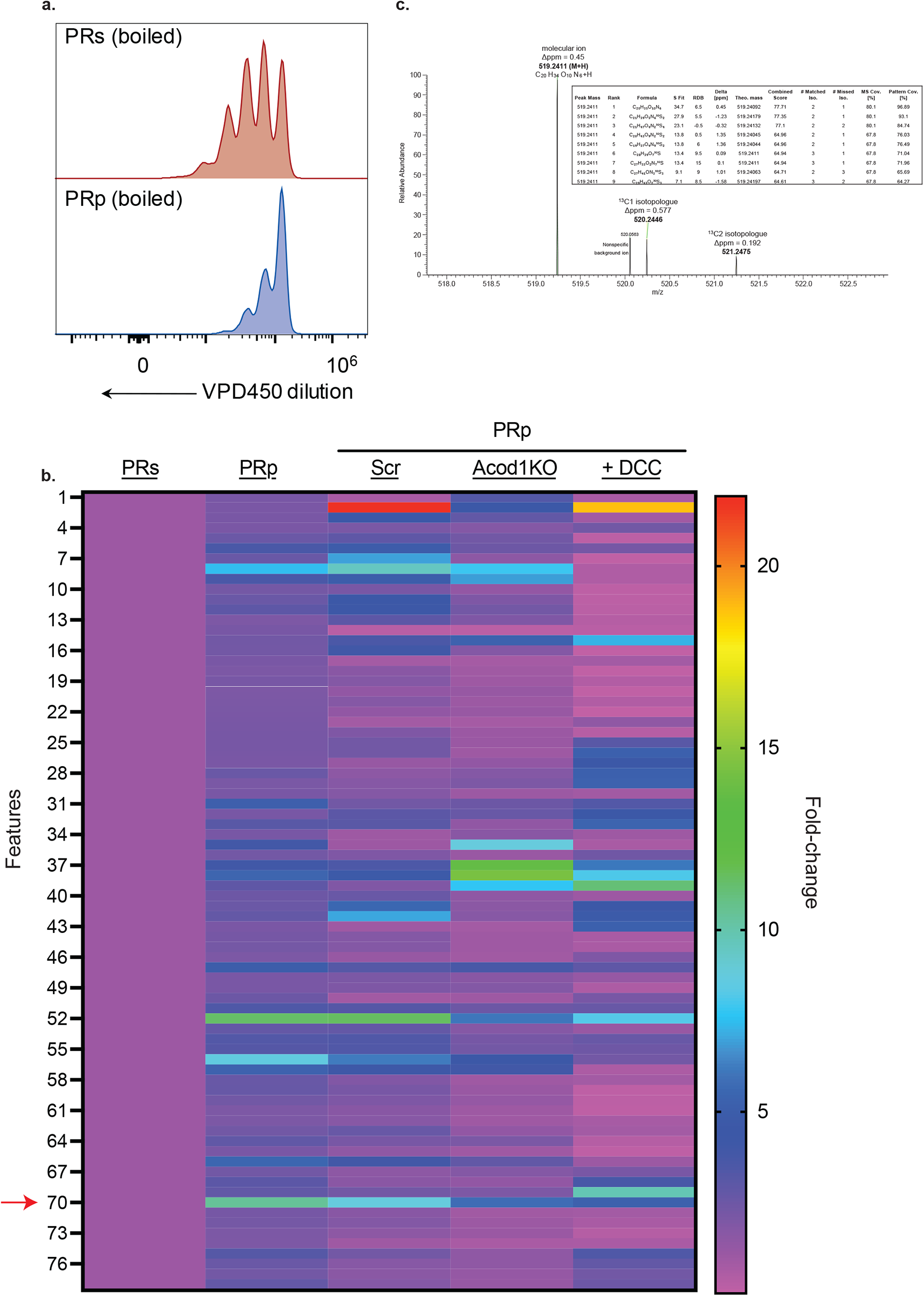
Identification of Acod1-mediated inhibitory peptides. **a.** Representative histograms of violet proliferation dye 450 (VPD450) dilution in CD8^+^ T cells following 72 h of activation with αCD3/CD28 in indicated boiled CM **b.** Heatmap depicting compounds detected by LC/MS in the indicated conditioned medium (CM) by fold-change relative to PRs CM of that compound. Each square represents the average of three replicates of that condition. (*n* = 3). **c.** MS1 peaks of m/z = 519.2411 compound.

